# Surface architecture of the bacterial envelope determines phage adsorption route in pathogenic *Escherichia coli* O157:H7

**DOI:** 10.64898/2026.07.23.740414

**Authors:** Edwin Omar Rivera-Lopez, Lucas Morinière, Alexey Kazakov, Denish Piya, Melina Pena, Benjamin Weinberg, Jie Feng, Avery Noonan, David W. Lacher, Adam M. Deutschbauer, Adam P. Arkin, Vivek K. Mutalik, Edward G. Dudley

## Abstract

The outermost surface layers of Gram-negative bacteria determine phage access to terminal receptors, yet their genetic basis has been mapped almost exclusively in laboratory strains that lack them. Here we apply genome-wide RB-TnSeq fitness profiling to four *Escherichia coli* O157:H7 strains from distinct phylogenetic clades sharing the O157 O-antigen, using 38 phages with terminal receptors previously mapped in *E. coli* K-12 strain. RB-TnSeq fitness landscapes across all four pathogenic backgrounds were mostly similar, and dominated by surface-associated loci, including the *gfc-etk* group 4 capsule operon, O-antigen biosynthesis genes, LPS core assembly genes and outer membrane proteins. Disruption of *gfc-etk* abolished infection in 11 genetically diverse myoviruses, establishing the O-antigen capsule as a widespread required primary recognition substrate. O-antigen loci generated two classes of fitness score patterns. For 10 phages, disruption increased infectivity, indicating it is a barrier to receptor access; for 3 others, disruption abolished infectivity, demonstrating it can also be a primary recognition substrate. Outer membrane protein receptor identity was conserved across laboratory and pathogenic backgrounds, with the same proteins recognized in both K-12 and O157:H7, while glycan layer state determines whether these receptors are reached. These results demonstrate that outer surface glycan layers can act as primary and optional recognition substrates for phage infection, or as physical barriers preventing terminal receptor access. Extending the ability to probe phage-targeted receptors beyond outer membrane proteins provides a framework for incorporating glycan layer state into predictive models of phage-host interactions.

**Importance:** Bacteriophage-based interventions for controlling *Escherichia coli* O157:H7, a major foodborne pathogen responsible for tens of thousands of illnesses annually in the United States, require a mechanistic understanding of the factors governing strain-level susceptibility. Predictive frameworks developed in laboratory model strains lacking O-antigen and capsular polysaccharides can map the terminal protein receptors that phages bind, but are currently limited in their ability to determine whether those receptors are accessible in pathogenic isolates carrying full outer surface complexity. This study provides the first genome-scale, functional genetic map of phage susceptibility determinants in O157:H7 and demonstrates that the state of the outer surface layers, specifically the O-antigen and the *gfc-etk* capsule, determines whether phages can reach conserved terminal receptors. This finding explains differences in phage susceptibility between strains sharing nearly identical gene content, and identifies the molecular layers that must be characterized to predict phage host interaction in pathogenic *E. coli* backgrounds.

## Introduction

*Escherichia coli* O157:H7 is among the most intensively studied foodborne pathogens, responsible for an estimated 86,000 illnesses annually in the United States and accounting for the majority of Shiga toxin-producing *E. coli* outbreaks linked to contaminated beef and fresh produce (1–3). The serotype encompasses multiple genetically distinct lineages distributed across at least nine phylogenetic clades defined by single-nucleotide polymorphism analysis (4). These lineages differ in virulence potential and prophage content, yet they share the O157 O-antigen and additional surface-associated structures that distinguish them from laboratory model strains and from other *E. coli* pathotypes.

Phage-based approaches to controlling O157:H7 in food production and clinical settings have attracted sustained interest (5, 6). The feasibility of these approaches depends on identifying phages capable of infecting the relevant strain diversity, which requires an understanding of what makes particular strains susceptible. Historically, phage susceptibility across O157:H7 isolates was captured through typing schemes that demonstrated reproducible strain-level differentiation based on lytic patterns, with epidemiologically linked isolates displaying similar phage types and unrelated isolates showing substantial variability (7, 8). These observations established that phage susceptibility is a strain-level phenotype within O157:H7, but the genetic and molecular basis of this variation has not been systematically resolved.

Mechanistically, phage infection of Gram-negative bacteria proceeds through a sequential adsorption process. Initial reversible interactions with surface glycans, including O-antigen polysaccharide (OPS) or capsular exopolysaccharides (CPS), bring the phage into proximity with the outer membrane. Irreversible binding to a terminal receptor, typically an outer membrane protein (OMP) or a deep-core LPS sugar, then commits the phage to genome injection (9, 10). In laboratory strains such as *E. coli* K-12 and its derivatives, including BW25113, which lack functional O-antigen polysaccharide and capsular polysaccharides (11), only the terminal receptor engagement step is operative and phages interact directly with outer membrane proteins and LPS core sugars without encountering upstream glycan layers. This property allowed Moriniere et al. (12) to map terminal receptor specificity systematically for 255 double-stranded DNA (dsDNA) phages across 19 receptor classes using genome-wide RB-TnSeq (random bar code transposon site sequencing (13)) and Dub-seq (Dual-barcoded shotgun expression library sequencing (14)) screens in BW25113, establishing a comprehensive receptor reference for a large and taxonomically diverse phage collection.

Most clinically and environmentally relevant *E. coli* lineages, including O157:H7, express O-antigen polysaccharide and in some cases additional capsular exopolysaccharides that form the outermost layers of the cell envelope (**Figure 1**, sugar chain composition derived from literature sources (15–18)). Specifically, *E. coli* O157:H7 and other Shiga toxin producing *E. coli* lineages (STEC) possess a group 4 capsule (G4C) (19, 20), also referred to as the O-antigen capsule as both polysaccharides are made of the same tetrasaccharide repeat units (16), whose export is encoded in the *gfc*-*etk* operon (21). These structures are not passive bystanders in phage-host interactions. The O-antigen can shield outer membrane protein receptors from phage receptor-binding proteins, requiring phages to process or bypass the glycan layer to reach their terminal site. Alternatively, specific phages engage O-antigen directly as a primary recognition substrate, with this interaction being essential for subsequent adsorption (22, 23). The G4C was also recently demonstrated to be the primary receptor of a T4-like phage in the O157:H7 model strain EDL933, while multiple LPS core sugars were also recognized as secondary receptors (24). The biosynthetic gene cluster encoding the O157 antigen is conserved in its genetic organization across STEC isolates (25), and *E. coli* O-antigen gene clusters more broadly have been systematically catalogued into biosynthetically related groups sharing nearly identical genetic architectures (26). This conservation means that the genomic presence of O-antigen biosynthesis genes provides limited resolution on the surface state differences that determine phage access across strains within a serotype.

**Figure 1.**
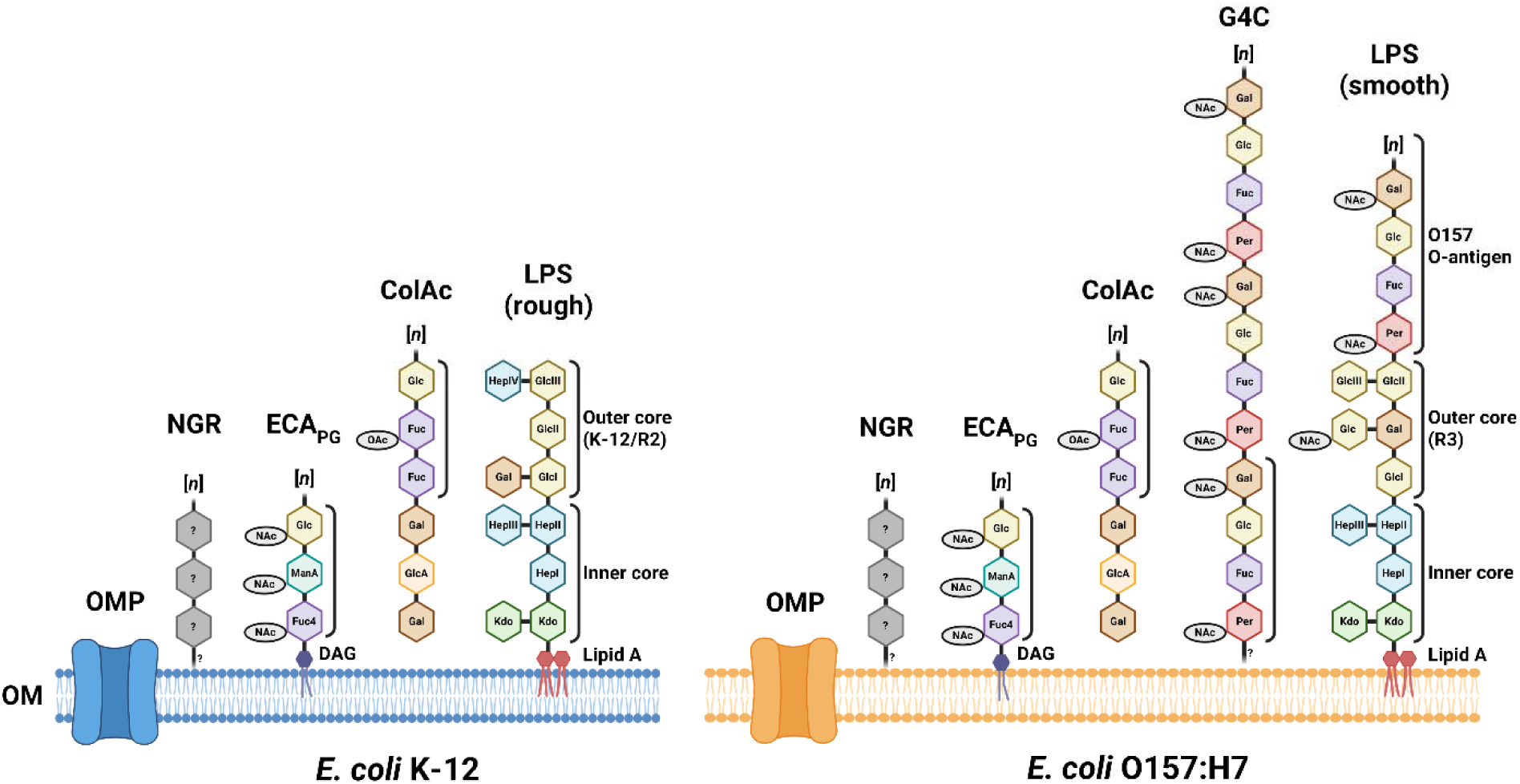
Schematic of the outer surface glycan composition and content of *E. coli* K-12 and O157:H7. Polysaccharide chain composition was derived from literature (Miravet-Verde et al. 2026, Sande and Whitefield 2021, Wang and Reeves 1998, Kai and Mitchell 2020). Presence of a R3-type LPS outer core in O157:H7 strains was deduced from LPS biosynthetic gene cluster organization (Miravet-Verde et al. 2026). Enterobacterial common antigen is displayed only in its most abundant peptidoglycan-linked form (**ECA_PG_**). Brackets indicated tri and tetrasaccharide repeat units. **OM** = Outer membrane; **OMP** = Outer membrane protein; **LPS** = Lipopolysaccharide; **G4C** = Group 4 capsule; **ColAc** = Colanic acid capsule; **NGR** = N4-glycan receptor; **Kdo** = 3-deoxy-D-manno-octulosonic acid; **Hep** = Heptose; **Glc** = Glucose; **Gal** = Galactose; **Per** = Perosamine; **Fuc** = Fucose; **GlcA** = Glucuronic acid; **Fuc4** = 4-acetamido-4,6-dideoxy-D-galactose; **ManA** = Mannosaminuronic acid; **DAG** = Diacylglycerol; **NAc** = N-acetyl group; **OAc** = O-acetyl group. Figure created in Biorender.

To determine what governs phage infectivity across O157:H7 strain backgrounds, we performed genome-wide RB-TnSeq fitness profiling in four representative O157:H7 strains using a subset of the phage collection characterized in BW25113 by Moriniere et al. (12). This experimental design situates the study to resolve which host factors, beyond terminal receptor identity, control phage access in pathogenic backgrounds carrying full outer surface complexity. The results presented here identify the state of the outer surface architecture, specifically the O-antigen and the *gfc-etk* capsule, as a major determinant of phage infectivity across O157:H7 strain backgrounds and establish a genome-scale functional framework for interpreting phage-host interactions in clinically relevant isolates.

## Results

### Bacterial strain selection and genomic characterization

Four O157:H7 isolates representing distinct phylogenetic clades were selected for this study: ECRC98 (clade 7.2), ECRC100 (clade 6), ECRC101 (clade 8.3), and ECRC102 (clade 2) (4) (**Figure 2A**). These strains were initially chosen to sample the breadth of O157:H7 phylogenetic diversity and test genetic backgrounds with different predicted antiviral defense system (DS) repertoires. Pangenome analysis identified 4,745 gene families present in all four strains alongside 580 accessory gene families distributed variably across the panel (**Figure S1**). ECRC98 and ECRC100 showed the highest pairwise gene family overlap, reflecting close genomic similarity.

**Figure 2.**
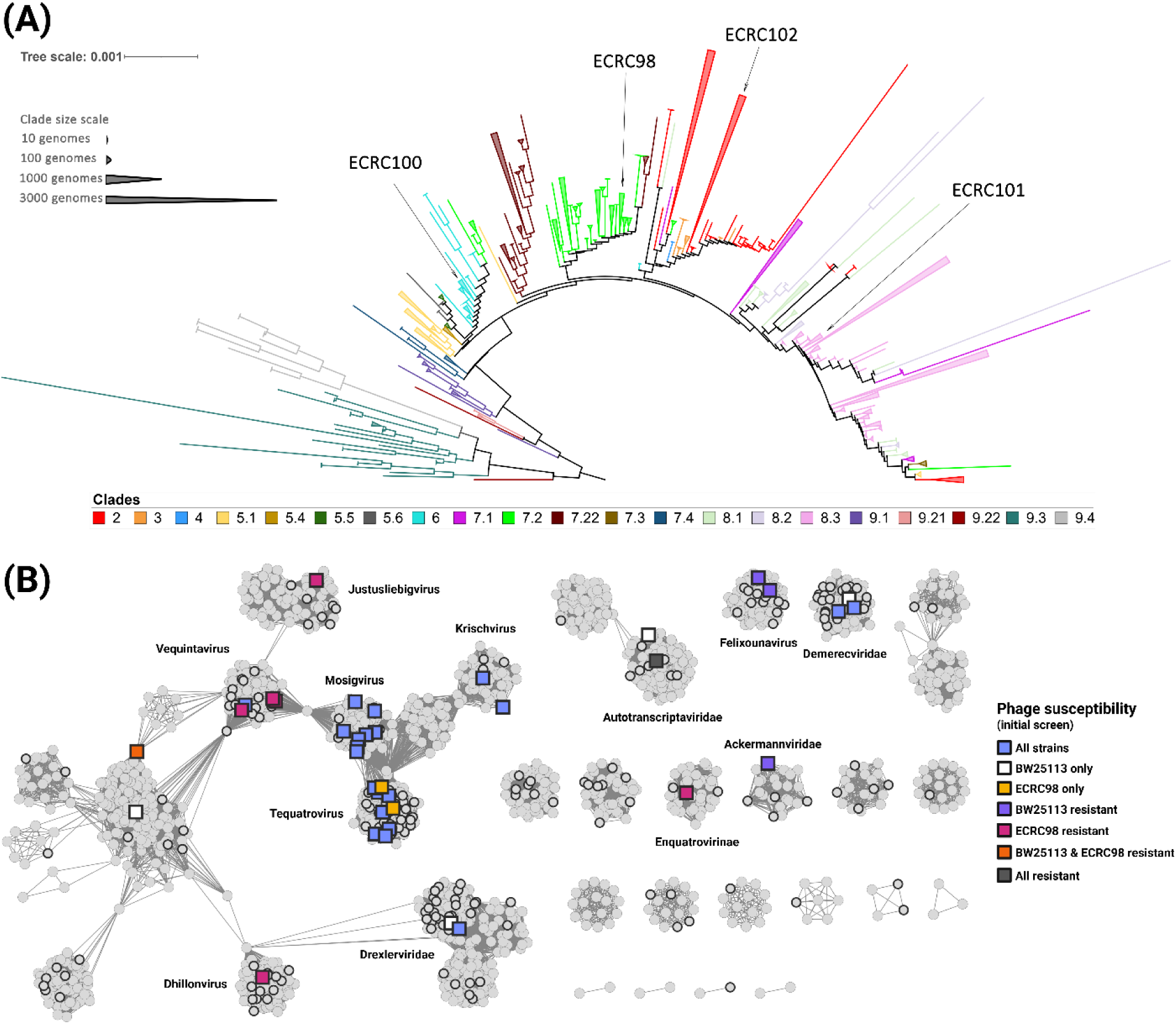
Biological resources used in this study. **(A)** Whole genome-based phylogeny of 32,252 genome-sequenced *E. coli* O157:H7 isolates showing the positions of the four selected strains (ECRC98, ECRC100, ECRC101, and ECRC102). Known O157:H7 clades are indicated by different colors. Clades 9.3 and 9.4 correspond to *E. coli* O55:H7 lineages. **(B)** vConTACT2 gene-sharing network of 1,875 *E. coli* dsDNA phage genomes retrieved from the NCBI GenBank database. Isolates from the collection assembled by Moriniere et al. (2026) are indicated with a black outline. The 38 phages selected for this study are indicated by a square shape, and colors represent their ability to induce bacterial clearance on the 5 strains tested during the initial screen (BW25113, ECRC98, ECRC100, ECRC101, and ECRC102).

Outer surface glycan biosynthetic gene clusters, including O157 O-antigen and lipopolysaccharide (LPS), group 4 capsule, colanic acid capsule, enterobacterial common antigen (ECA), and N4-glycan receptor (NGR), as well as phage-targeted outer membrane protein receptor genes (*fadL*, *tsx*, *ompA*, *ompF*, *ompW*, *ompC*, *fhuA*, *btuB*, *lptD*, *yncD*, and *lamB*) were compared between strains to search for non-synonymous single nucleotide polymorphism (SNPs) susceptible to interfere with phage recognition. A nonsense mutation was detected within the ORF of *nfrA* in the genome of ECRC98, preventing the secretion of the NGR polysaccharide in this strain (**Table S1**). Outer surface composition was otherwise genetically identical across the 4 O157:H7 backgrounds.

To assess whether variation in antiviral defense gene content could also affect phage susceptibility across the panel, and more broadly across the O157:H7 lineage, we annotated predicted defense systems across 32,252 O157:H7 genomes, including the 4 selected strains, using PADLOC (27) and DefenseFinder (28). Core systems including restriction-modification (RM) types I, II, and III, CRISPR-Cas class 1 subtype I-E, and MazEF toxin-antitoxin were present in over 99% of genomes with no significant clade-specific enrichment or depletion (**Note S1, Figure S2, Datasets S1-S2**). Across the 4-strain panel, 42 genes predicted to be collectively encoding 21 distinct DS were shared in all backgrounds, and a common Lamassu-family defense system (29) was predicted in every genome except for ECRC101 (**Table S2**). All backgrounds also encoded strain-specific defense systems: ECRC98 contained a unique type II RM system, an SDIC3 system (30), and 2 phage defense candidate systems (PDC-S39, PDC-S25); ECRC100 contained unique RosmerTA (29), DarTG (31), Zorya (32), and type I RM systems; ERC101 contained a distinct Lamassu-family DS from the one found in other strains, an abortive infection system (PD-T4-6) (33) and 2 phage defense candidate systems (PDC-S43, PDC-M11); finally, ECRC102 only contained a unique type II RM system. Defense system composition is therefore primarily conserved across O157:H7 phylogenetic diversity, although substantial combinatorial variation arises from how individual systems are assembled within genomes.

### Phage strain selection and infectivity across the O157:H7 strain panel

A panel of 38 phages was selected from the 255-phages collection characterized by Moriniere et al. (12). The primary selection criteria were coverage of diverse phage families and receptor classes, and visible bacterial clearance on some or all of the O157:H7 strains in an initial susceptibility screen where 161 phages from the collection were prepared at a high titer (10^9^ PFU/mL) and spot-tested on bacterial lawns (**Figure 2B, Dataset S3**). Five phages which did not inhibit growth on any of the O157:H7 strains in plate assays were still included in the panel to increase coverage of phage taxa and receptor classes (Bas10, EV219, Lambda, Bas68, and K30), and to test whether genome-wide screens could identify genes for which disruption induces phage susceptibility.

A complementary susceptibility screen was conducted to assess whether phages in the panel could form visible single plaques on the O157:H7 strains, and estimate their efficiency of plating (EOP) compared to the bacterial hosts used for their amplification (MG1655ΔRM and other strains). Ten phages consistently produced clear single plaques at a high EOP in the 4 strains, while other phages failed to form visible single plaques on at least one strain for which clearance was recorded during the initial screen (**Table S3**). This initial growth inhibition phenotype was yet always reproduced in this second screen, at least at the highest phage titer and most often across lower dilutions, with the exception of phage RB49 for which discrepancy between the two screens was attributed to a lower titer of the phage lysate used in the second screen.

As expected, most phages using the NGR as their receptor proved unable to infect strain ECRC98 (phages Bas58, EV146, TP5, Bas61, and Bas69), in which secretion of the polysaccharide is impaired by the mutation in *nfrA* (34). However, the high infectivity on this strain of phage Bas48, which recognizes the NGR in a K-12 background, suggests that it can alternatively recognize another terminal receptor to complete its adsorption.

Although strains ECRC100, ECRC101 and ECRC102 displayed identical susceptibility profiles in the initial screen, notable differences were observed in the second assay, including dramatic reductions in EOP (> 10,000-fold) and inability to form visible single plaques for phages Bas48 and TP5 in ECRC102, phage TP9 in ECRC100, and phage TP6 in ECRC101. This tight phage-to-strain relationship (the pairwise proteomic equivalent quotient (35) between phages TP3 and TP6 is 0.975, yet only TP6 displayed this impaired phenotype), and the occurrence of bacterial clearance without single plaque formation, suggests the possible action of strain-specific defense systems in modulating phage infection post-adsorption.

In total, this 38-phages panel covered 16 phage genera and 14 receptor classes (**Table S3**). These phages were isolated on different bacterial hosts than the O157:H7 strains used in this study, although 11 phages were isolated using other O157:H7 backgrounds. It is thus unlikely that any of the phages in the subset have been previously exposed to the specific defense systems present in the 4 O157:H7 backgrounds tested. Receptor specificity could not be resolved by previous genome-wide screens in BW25113 for 4 of the O157:H7-specific phages (TP9, TP11, TP15, and CBA120). A functional genetic approach is therefore necessary to resolve the operative determinants of susceptibility for these phages, and assess whether other phages recognize outer surface glycan layers in addition to their terminal OMP receptors in O157:H7 pathogenic backgrounds.

### Genome-wide fitness profiling identifies outer surface architecture as the primary determinant of phage susceptibility

To identify the host genetic factors governing phage infectivity, we constructed RB-TnSeq transposon mutant libraries in all four O157:H7 backgrounds and challenged them with phages in liquid fitness assays following the protocol initially developed for phage receptor characterization in BW25113 (12). This design allows direct comparison of O157:H7 fitness landscapes against the BW25113 receptor reference established in that study. Fitness scores were calculated from barcode sequencing of surviving mutant populations relative to pre-infection reference samples, with high-confidence effects defined by a fitness score at or above 4 (log2 fold enrichment) and an absolute t-statistic at or above 5. RB-TnSeq fitness profiling detects genes whose disruption alters host survival under phage pressure and is structurally suited to identify adsorption-stage determinants. However, the strength of the selection (10:1 ratio of phage particles over bacterial cell) is such that secondary effect mutations unrelated to surface determinants which could still affect survival rate are masked by dominant mutations rendering cells impervious to phage infection. For phage-strain combinations in which infection proceeds efficiently in liquid culture, disruption of intracellular defense systems produces no detectable fitness difference under these assay conditions, as defense gene mutants are likely wiped out from the population early in the assay.

Fitness landscapes across the ECRC10, ECRC101 and ECRC102 backgrounds were dominated by genes encoding surface-exposed structures and are the primary basis for the results reported below (**Figure 3**, **Figures S3-S6**, complete gene lists in **Datasets S4**-**S11**). These fitness signals organized into four functional groups: group 4 capsule biosynthesis and export with the *gfc*-*etk* operon (phages LZ9, RB51, RB68, Bas38, K20, Bas47, CEV1, Bas48, TP5, TP11, TP15); O-antigen/ECA/NGR biosynthesis and modification genes including *gnu* (RB68, CBA120, TP9), *wzzB* (JK32), and *nfrA*/*nfrB* (Bas58 and Bas69, only in ECRC101 and ECRC102); LPS core assembly *waa* genes for every sugar residue above the first heptose in the inner core (diverse combinations for RB51, RB68, Bas38, K20, Bas47, CEV1, Bas48, TP5, JK32, CBA120, TP11, TP15); and outer membrane proteins *ompC* (Bas41, JK36, JK42, TP3, TP6), *ompF* (RB51, RB68, Bas38, Bas47, K20, CEV1), *ompA* (LZ9, RB49), *tsx* (JK32), *fadL* (T2), *fhuA* (CEV2), and *btuB* (EV212). This pattern was reproducible with minor variations across independently constructed libraries in ECRC100, ECRC101, and ECRC102. The ECRC98 library yielded broadly consistent results for outer membrane protein and group 4 capsule loci; however, genome-wide assays conducted with this library produced ambiguous fitness signals for O-antigen and LPS core biosynthesis genes, as mutants in these loci displayed elevated fitness in no-phage control experiments. A plausible explanation is that biosynthesis of O-antigen polysaccharide and LPS core sugars imposes a measurable metabolic cost and their disruption confers a modest growth advantage under the competitive broth culture conditions of the fitness assay. This background enrichment in the absence of phage selection renders fitness scores for these loci in ECRC98 uninterpretable and they are excluded from downstream comparisons involving O-antigen and LPS core assignments. Overall, these results confirm that surface-associated loci consistently represent the dominant fitness determinants during phage challenge across O157:H7 genetic backgrounds.

**Figure 3.**
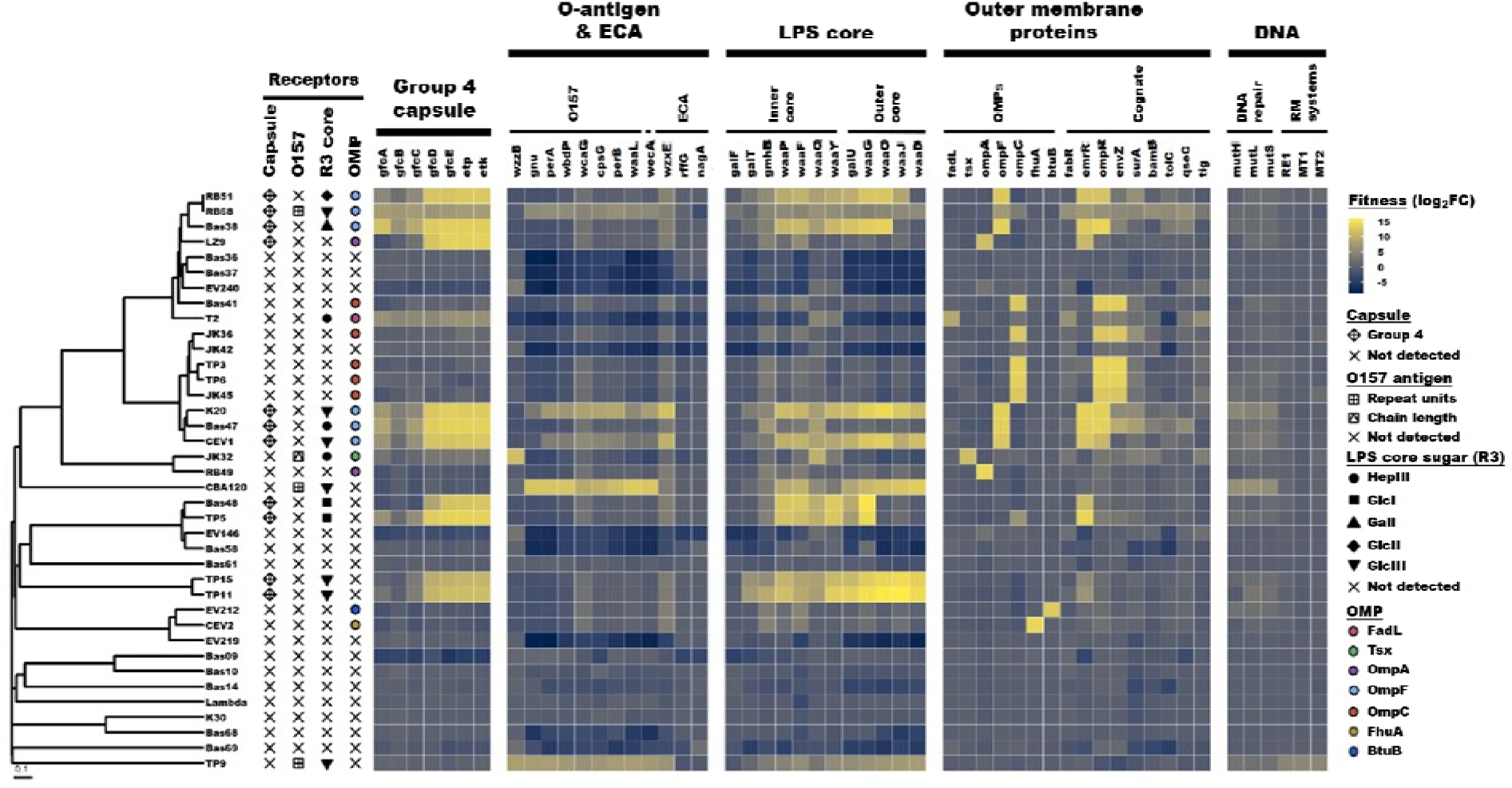
Genome-wide identification of surface-associated determinants of phage susceptibility in *Escherichia coli* O157:H7 strain ECRC100. Heatmap of loss-of-function (LOF) RB-TnSeq selected data for 38 dsDNA phages, where each row represents a phage and each column represents a gene. Fitness values are shown as log₂ enrichment scores, where high values indicate a selective advantage under phage challenge, and low values indicate a detrimental fitness effect. Genes are grouped by functional categories, including group 4 capsule, lipopolysaccharide (LPS) core, O157 O-antigen, outer membrane proteins (OMPs), and transport-associated functions. Other acronyms: RM = restriction-modification; RE = restriction endonuclease; MT = methyltransferase. Multi-gene signals were summarized into recognition of unique outer surface components (capsule, O157 O-antigen, R3-type LPS core sugars, OMPs). Phage dendrogram was computed based on pairwise proteomic equivalent quotient (PEQ) values.

A key feature of the fitness data is the presence of two mechanistically distinct classes of fitness signal for O-antigen loci. For a substantial number of phages (T2, Bas36, Bas37, EV240, JK42, Bas58, EV146, EV219, Bas68, and Bas69), disruption of O-antigen biosynthesis genes produced negative fitness scores in at least one library, indicating that mutants with disrupted O-antigen were susceptible to phage killing while cells displaying a functional O-antigen were impervious to phage infection. This result directly demonstrates that the intact O-antigen was restricting phage access to otherwise susceptible cells in those interactions: removing the glycan layer made infection possible. For a second smaller group of phages (RB68, CBA120, and TP9) the same gene disruptions produced positive fitness scores, indicating that O-antigen absence impairs phage infection. In phages TP9 and CBA120, for which only the mutants of O-antigen and full LPS core loci display high fitness scores, the O-antigen is engaged as a primary recognition substrate, and infection requires that engagement. The additional high fitness scores for group 4 capsule loci with phage RB68 suggests that the capsular polysaccharide could be the recognized factor instead of the O-antigen. Indeed, the tetrasaccharide repeat units composing these two polysaccharide chains are identical, and they share the same biosynthetic pathway (21). Disruption of O-antigen loci leads to the absence of both O-antigen and G4C, while disruption of G4C export does not remove O-antigen from the cell surface architecture. Disruption of the O-antigen chain length determining gene *wzzB* was sufficient to produce a strong positive fitness score in assays with phage JK32, indicating that variation in O-chain length alone reduces infectivity. Both classes of signal (negative and positive fitness scores) were detected within the same strain backgrounds with little to no differences between strains, establishing that the functional role of the O-antigen in a given interaction is a property of the phage and not of the strain.

The *gfc-etk* operon, encoding the group 4 capsule export system in O157:H7 and related STEC strains, emerged as high-confidence fitness loci for a subset of phages across multiple O157:H7 backgrounds (RB51, RB68, Bas38, LZ9, K20, Bas47, CEV1, Bas48, TP5, TP11, and TP15). Strong positive fitness scores indicated that capsule disruption impaired or abolished infection, showing that group 4 capsule is a required determinant for these phages. Prior characterization of the group 4 capsule in O157:H7 focused on its role in transiently shielding virulence-associated surface structures from extracellular recognition during host-cell contact (19). The fitness data presented here indicates that this capsule layer also participates in phage adsorption, likely functioning as a primary receptor and that these phages cannot interact nor bypass the O-antigen glycan layer when the capsule is disrupted. Inactivation of G4C export genes *etp* and *etk* was reportedly associated with an increased abundance of O-antigen, as the shared repeat units are redirected towards the O-antigen biosynthesis pathway (21), possibly explaining this phenomenon. Nonetheless, this represents the first such example indicating group 4 capsule as a phage-relevant surface structure in O157:H7.

In addition to identifying the O-antigen as the primary recognition substrate of phages CBA120 and TP9, unresolved by previous screens in BW25113, genome-wide screens in O157:H7 backgrounds similarly revealed that both the G4C and the last glucose of the R3-type LPS outer core (GlcIII) were the primary determinant of susceptibility for *Felixounavirus* phages TP11 and TP15, as indicated by high fitness scores for all the *waa* biosynthetic genes, including *waaD* which mediates the addition of the GlcIII to the outer core. Extending the functional genetic screening strategy to these pathogenic backgrounds thus successfully identified the operative determinants of susceptibility for all 4 O157:H7-specific phages that could not be resolved by screens in K-12.

### Functional validation of RB-TnSeq fitness genes

To confirm that the fitness signals identified by RB-TnSeq reflect direct contributions to phage susceptibility, we isolated individual transposon mutants from the ECRC100 and ECRC101 libraries in representative genes from each surface layer category: *etk* and *gfcE* from the capsule locus, *gnu* and *wecA* from the O-antigen pathway, *waaD* from the LPS core, and *tsx* and *ompA* from the outer membrane protein group. Complete results are reported in **Dataset S12**.

Efficiency of plating assays mostly confirmed that disruption of each gene altered susceptibility to the predicted phage or phages in a direction consistent with the corresponding RB-TnSeq fitness score (**Figure 4**). For complementation, constructs driving gene expression from the constitutive ompA promoter of ECRC100 were used in place of native promoters to ensure consistent expression across all tested loci. This approach restored wild-type susceptibility fully or substantially for the majority of phage-gene combinations tested, confirming that the observed phenotypes were attributable to loss of the specific gene rather than to secondary polar effects. In cases where complementation did not fully restore susceptibility, including phage Bas38 on the *etk* mutant, the residual phenotype likely reflects differences between the constitutive promoter context and native expression levels rather than a failure of complementation per se. Among the validated loci, disruption of *etk* in ECRC100 (**Figures 4A-B**) and *gfcE* in ECRC101 produced the largest decreases in susceptibility and most often completely abolished phage infection, consistent with the group 4 capsule being a primary recognition substrate required for phage access to its terminal receptor in these backgrounds. These assays also resolved weak and ambiguous RB-TnSeq fitness scores for a subset of phage-gene pairs, showing that O-antigen loci in phages K20 and CEV1, and group 4 capsule loci in phages T2 and JK32, do not contribute to phage susceptibility and are excluded from receptor assignments.

**Figure 4.**
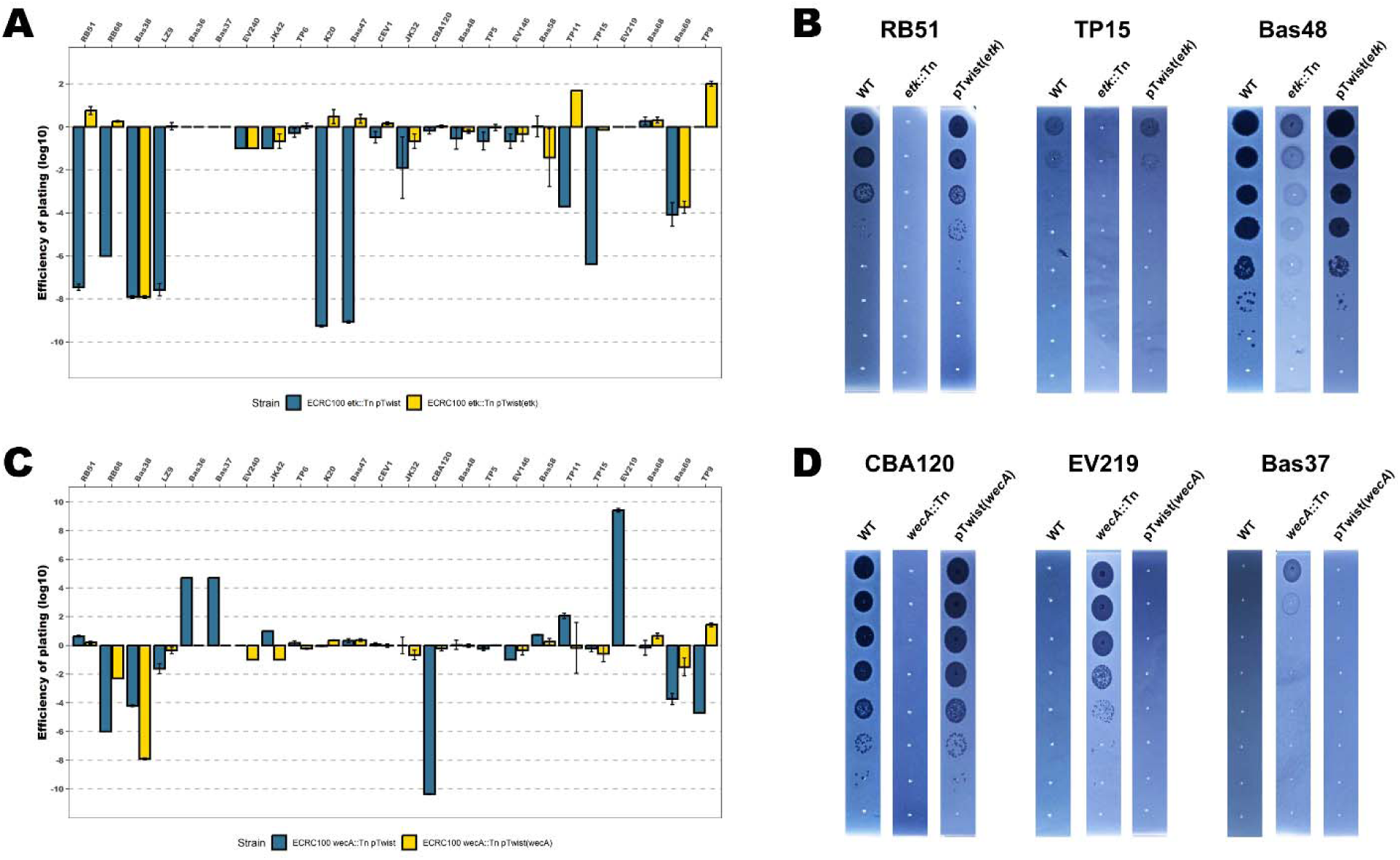
Validation of RB-TnSeq fitness signals with 24 selected phages on single-gene transposon mutants in ECRC100. (A-B) Efficiencies of plating on an ECRC100 *etk*::Tn capsular mutant, with or without complementation *in trans*. (C-D) Efficiencies of plating on an ECRC100 *wecA*::Tn O-antigen mutant, with or without complementation *in trans*. Experiments were systematically conducted in biological triplicates, and error bars indicate standard deviation from the mean. Absence of error bars indicate no variation was recorded between triplicates, and is often indicative of absence of single plaque formation. Panels 4B and 4D show representative instances where gene disruption abolishes infection (positive fitness class in RB-TnSeq) or enables infection (negative fitness class in RB-TnSeq). Bas48 on ECRC100 *etk*::Tn is shown as an example of cases where capsule disruption impairs plaque phenotype without quantitatively altering infection efficiency.

Similarly, disruption of O-antigen loci *wecA* in ECRC100 (**Figures 4C-D**) and *gnu* in ECRC100 validated both positive and negative fitness scores. Infection was abolished or severely impaired for phages RB68, TP9 and CBA120, confirming the O157 O-antigen is a required substrate reversibly bound and/or cleaved upon recognition, as it was shown previously for CBA120 (Plattner et al. 2019). On the other hand, some phages that were not showing signs of susceptibility on the wild-type host (Bas36, Bas37, EV219, and Bas68) gained the ability to efficiently infect the mutant strains, proving that the O157 O-antigen can solely modulate phage susceptibility by blocking access to the phages’ terminal receptors.

### Outer membrane protein recognition substrate identity is conserved across host backgrounds while adsorption route is not

To determine the relationship between phage adsorption determinants in O157:H7 and the receptor assignments established in BW25113, we compared RB-TnSeq fitness data from all four O157:H7 libraries to the BW25113 data from Moriniere et al. (12). We found that OMP terminal receptor identity is conserved for every phage but one across K-12 and O157:H7 host backgrounds: OmpF (RB51, Bas38, K20, Bas47, and CEV1), OmpC (Bas41, JK36, JK42, JK45, TP3, and TP6), OmpA (LZ9, EV240, and RB49), Tsx (JK32, Bas36 and Bas37), FadL (T2), FhuA (CEV2), and BtuB (EV212). Phage RB68 constituted the only exception to this rule, with OmpF being recognized in every O157:H7 background in place of OmpW in BW25113. We previously reported a similar OmpF-specific phenotype in BL21, and it should be noted that targeting of OmpW in BW25113 results from a single point mutation at the binding interface of the phage’s RBP. These results demonstrate that OMP terminal receptor identity is therefore an intrinsic property of the phage that is preserved regardless of whether the host carries an outer glycan layer. OMP receptor assignments from BW25113-based screens are transferable to pathogenic backgrounds for the terminal receptor step of adsorption, although it does not predict whether adsorption will succeed in a pathogenic background, because that outcome depends on whether and how the phage interacts with the glycan layers that gate receptor access.

Several groups of phages illustrate this at the level of individual phage-strain comparisons. High fitness scores for group 4 capsule loci in *Tevenvirinae* phages recognizing OmpF as their primary receptor indicates that these phages engage a surface component in specific O157:H7 backgrounds that does not exist in BW25113. The theoretical 3-step recognition sequence operated by these phages during adsorption to O157:H7 outer membrane therefore begins with specific binding to the group 4 capsule using an unknown factor, followed by reversible binding of the Gp38 adhesin to OmpF, and ends with the irreversible attachment of tailspikes Gp12 to the first heptose of the LPS inner core, triggering contraction of the tail tube and DNA injection (9).

Phages in the *Vequintavirus* (Bas48, Bas58, TP5, EV146), *Justusliebigvirus* (Bas61), and *Enquatrovirus* (Bas69) taxa constitute other interesting examples of how terminal receptor identity is conserved while prior interaction with glycan layers can vary across strain backgrounds. RB-TnSeq screens conducted in BW25113 with 18 *Vequintavirus* and 8 *Justusliebigvirus* phages led to the conclusion that they were all targeting the NGR polysaccharide as their primary recognition substrate, and the first glucose of the LPS outer core (GlcI) as their terminal receptor. *Enquatrovirus* phages Bas69 and N4 were relying solely on the NGR to complete their adsorption (10, 12, 34). The picture is actually much more complex in O157:H7 backgrounds: Bas48 and TP5 still use the GlcI sugar, likely as their terminal receptor, but interact with the group 4 capsule instead of the NGR in ECRC100, ECRC101, and ECRC102. On the other hand, Bas58 and EV146 both yielded negative fitness scores for the O-antigen loci in ECRC100, and gave a signal consistent with recognition of both the NGR polysaccharide and the GlcI sugar in ECRC101 and ECRC102, although the signal was very weak for EV146 as infection did not seem to proceed efficiently in liquid assays. Although it shared the exact same combination of recognition substrates in BW25113, *Justusliebigvirus* phage Bas61 did not yield any significant fitness score in these strains and seemed unable to infect them. Finally, Bas69 also displayed negative fitness scores for O-antigen loci in ECRC100, and positive scores for only the NGR loci in ECRC101 and ECRC102.

These results indicate that recognition steps upstream of terminal receptor binding actually differ between *Vequintavirus* phages and across bacterial hosts. Although NGR biosynthetic loci are identical across O157:H7 strains, it seems that the NGR polysaccharide is not accessible to phage and possibly masked by the O-antigen in the specific ECRC100 background. The unique property of Bas48 and TP5 to recognize the G4C instead of the NGR in the ECRC100 background likely reflects the presence in their genome of a multivalent set of alternative tail fibers able to recognize different polysaccharide substrates depending on the host background (Schwarzer et al. 2012). In summary, the terminal receptor is the same across all backgrounds, while the upstream surface engagement differs by strain. Together these examples demonstrate that the adsorption route taken by a phage to reach its terminal receptor can differ across O157:H7 strain backgrounds that otherwise carry equivalent receptor gene content.

## Discussion

This study provides the first genome-scale functional genetic characterization of phage susceptibility determinants across multiple phylogenetically distinct *E. coli* O157:H7 strain backgrounds. By combining defense system analysis across 32,252 O157:H7 genomes, phenotypic profiling against a taxonomically diverse phage collection, RB-TnSeq fitness screening in four O157:H7 backgrounds, and direct comparison with receptor data from the laboratory strain BW25113, we identify three interconnected findings. First, predicted antiviral defense systems are highly conserved across O157:H7 clades and cannot explain the strain-level susceptibility differences observed in phenotypic assays. Second, the dominant fitness determinants in all four O157:H7 backgrounds are surface-associated loci, including group 4 capsule, O157 O-antigen, R3-type LPS core, ECA/NGR polysaccharide, and outer membrane proteins, with O-antigen loci generating two functionally distinct classes of fitness signal reflecting its role as a barrier to phage access and as a primary recognition substrate depending on the phage. Third, OMP receptor identity is conserved between BW25113 and O157:H7 backgrounds, while what differs between a simplified laboratory strain and a pathogenic isolate is the set of glycan layers the phage must navigate before reaching the terminal receptor that determines genome injection.

The dual role of O-antigen in phage adsorption has been established in individual model systems but has not previously been resolved at the scale of a diverse phage panel within a single pathogenic serotype. In a short commentary, Broeker and Barbirz (22) summarized evidence pointing towards direct O-antigen engagement by tailspike proteins of the podovirus coliphage G7C being the irreversible binding step that initiates infection, repositioning the O-antigen from a physical obstacle to an essential receptor for O-antigen-specific phages. Letarov (23) synthesized the complementary perspective in a comprehensive review: for the majority of *E. coli* O-antigen types, the O-polysaccharide layer provides robust shielding of outer membrane proteins and represents an omnipresent factor shaping phage-host ecology and co-evolution. Within *E. coli* O157:H7 specifically, the biosynthetic gene cluster encoding the O157 antigen is conserved in genetic organization across STEC isolates (25), and the broader *E. coli* O-antigen biosynthesis gene cluster diversity places related serotypes in groups sharing nearly identical genetic architectures despite antigenically distinct structures (26). This genomic conservation means that comparative gene content analysis cannot resolve the surface layer variation that our functional data detect. Our results show that specific recognition of the O157 O-antigen is not a common trait across the 38 phages characterized: only 3 phages belonging to unrelated taxa (*Tequatrovirus*, *Kuttervirus*, and *Uetakvirus*) yielded positive O-antigen fitness signals consistent with the O-antigen being a primary recognition substrate for infection. However, negative O-antigen loci fitness signals indicate that shielding of host terminal receptor by the O-antigen can prevent or impair infection across a wide diversity of phage groups (*Tequatrovirus*, *Vequintavirus*, *Tequintavirus*, *Berlinvirus*, and *Enquatrovirus*). Humolli et al. (36) demonstrated that engineering K-12 to express O-antigen revealed entire phage genera not recoverable from standard K-12-based screens, indicating that required O-antigen recognition by phage is dominantly a phylogenetically-stratified trait limited to specific phage taxa. The fitness data presented here extend these observations by showing that within the O157 serotype, both barrier and receptor functions of the O-antigen are simultaneously operative for different phages challenging the same host background, and that which function predominates reflects phage recognition machinery rather than host genotype.

The conservation of outer membrane protein receptor identity across BW25113 and O157:H7 backgrounds connects this study directly to the systematic receptor mapping of Moriniere et al. (12). That work assigned terminal receptors to 193 phages across 19 receptor classes using genome-wide screens in BW25113 and BL21, both of which lack O-antigen polysaccharide and capsular polysaccharides. The receptor assignments generated in that surface-simplified system are confirmed here in O157:H7: the same outer membrane proteins serve as terminal receptors for a given phage regardless of the host surface complexity. This finding also provides a mechanistic interpretation for the ceiling in genomic prediction of phage-host interactions reported by Gaborieau et al. (37), who achieved an area under the receiver operating characteristic curve of 86% for predicting infection outcomes across the *Escherichia* genus using genomic features as predictors. The interactions that remain unexplained at that ceiling likely include cases where the glycan layer state in the host strain, which is not captured by gene presence in a genome assembly, determines whether the phage can reach a terminal receptor that the genome suggests should be accessible. Gene content identifies which surface structures can potentially be produced; it does not specify whether those structures are assembled in a physical form that a given phage can engage or is blocked by. Chain length regulation, post-assembly modification, and production level are all determined by regulatory and biosynthetic processes that may vary between strains with equivalent gene content. The ECRC100, ECRC101, and ECRC102 comparison is the empirical demonstration of this gap at the strain level within O157:H7.

The O157 O-antigen capsule finding extends the analytical framework to a surface layer in O157:H7 that has not previously been systematically characterized in the context of phage adsorption. The fitness data presented here show that *gfc-etk* disruptions alter phage susceptibility in the same subset of phages across all the strains tested, indicating that this group 4 capsule layer participates in phage-host interactions as an outer envelope structure above the O-antigen, and that its recognition is an inherent property of the phage. Wang et al. (24) recently reported two *Tequatrovirus* phages which infect O157:H7 through recognition of specific receptors: PSD2001 recognizes both the G4C and the R3-type LPS outer core sugars GalI, GlcII, and GlcIII, while PNJ212 targets OmpC directly. PSD2001 is a T6-like *Tequatrovirus* and its genome encodes a Gp38 adhesin, indicating that it must also recognize an unknown outer membrane protein (38). Our findings indicate that O157 G4C capsule-recognition is a common accessory trait among T-even phages, yet restricted to T6-like phages, as all the T4-like phages in our set directly recognized their OmpC receptor. Outer membrane protein receptor specificity seems otherwise decoupled from G4C specificity, as phages binding OmpF (6) and OmpA (1) were found to also recognize G4C. We also found the same R3-type LPS core sugars reported by Wang et al. to be receptors of a subset of T-even phages recognizing G4C and OmpF. These findings support the hypothesis of a 3-step adsorption model for these phages, where the group 4 capsule, OmpF, and LPS core sugars are sequentially recognized as the phage gets closer to the outer membrane. We also report that G4C recognition is not restricted to T-even phages, and is also an accessory trait in other myovirus phage taxa: *Vequintavirus* phages Bas48 and TP5 can use G4C as an alternative recognition substrate when the NGR polysaccharide is not accessible, and *Felixounavirus* phages TP11 and TP15 recognize the G4C prior to interacting with the GlcIII LPS core sugar. A role for the group 4 capsule in modulating phage access has not previously been systematically described, and the phages affected by this layer represent candidates for future structural and mechanistic characterization of how the outermost surface layer is navigated or required.

Taken together, the O-antigen and *gfc-etk* capsule data define the outer surface of O157:H7 as a layered filtering system in which the state of each component, and the capacity of each phage to engage or bypass it, jointly determine infection outcome. These layers are present in the pathogen and absent from the laboratory strains in which most phage receptor biology has been systematically characterized, which is why O157:H7 phage receptor specificity was not fully predictable from prior work in K-12 derivatives alone. The genome-scale functional map provided here establishes a foundation for incorporating surface architecture into predictive models of phage host range in pathogenic *E. coli* and more broadly in Gram-negative pathogens that express antigenically diverse glycan outer layers.

## Materials and Methods

### Bacterial strains, plasmids, primers and growth conditions

Bacterial strains, plasmids, and primers used in this study are listed in **Tables S4, S5** and **S6**, respectively. Bacterial cultures were grown at 37°C in LB broth with agitation at 200 rpm, or on LB agar plates. Unless specified otherwise, antibiotics were supplemented at a final concentration of 25 µg/mL for kanamycin (Kan) and 30 µg/mL for chloramphenicol (Cam), and calcium chloride (CaCl_2_) was systematically added at a final concentration of 10 mM in phage experiments. Bacterial strains were kept for long-term storage at –80°C in 15 % (v/v) glycerol. Plasmids, primers, and genomic DNAs were stored at –20°C. Plasmid were ordered from Twist Bioscience, enzymes were acquired from New England Biolabs (NEB), and primers were synthesized at Elim Biopharmaceuticals and Integrated DNA Technologies (IDT).

### Bacteriophage strains and propagation

Bacteriophage strains are listed in **Table S7** with their genome accession numbers and corresponding propagation hosts. Phage lysates were prepared following the standard procedures described in Moriniere et al. (12). Briefly, overnight bacterial cultures were diluted at an OD_600_ of 0.1 in LB broth supplemented with CaCl_2_. Cultures were incubated after phage infection for a maximum of 4 h or until complete bacterial clearance was observed, centrifuged and filter-sterilized on 0.22 µm to remove bacterial cells and debris, and lysates were routinely stored at 4°C. Phage titers were determined by spotting 2 µL droplets of 10-fold serial dilutions made in SM buffer (Teknova) supplemented with CaCl_2_ on bacterial lawns prepared from overnight cultures grown in 0.5% T-top agar following a standard double-layer agar procedure. High-throughput spotting was conducted using a MicroPro 20 semi-automated pipettor instrument (Mettler-Toledo Rainin). Experiments were systematically conducted in biological triplicate. Plaque formation was recorded after an overnight incubation at 37°C.

### Bacterial long-read genome sequencing

Genomic DNA of *E. coli* O157:H7 strains ECRC98, ECRC100, ECRC101, and ECRC102 were extracted from 1 mL overnight cultures using the Wizard® Genomic DNA Purification Kit (Promega Corporation) following manufacturer protocol. Long-read genome sequencing was carried out by Plasmidsaurus using Oxford Nanopore Technology with custom analysis and annotation. Genome sequences were deposited on NCBI Genbank under BioProject number PRJNA1458855, and genome and single-read archive (SRA) accession numbers are listed in **Table S4**.

### Comparative genomic analysis of O157:H7

The *E. coli* O157:H7 genomes and clade designations were assigned using SNP-based identification (**Dataset S13**) (4). A total of 32,252 O157:H7 genome assemblies were retrieved from NCBI in GenBank format using the Bio.Entrez module of Biopython. Assemblies lacking annotated genes were re-annotated with Prokka v1.14.6 (39) using Prodigal (40) for protein-coding region prediction.

Phage defense systems were predicted using PADLOC v2.0.0 (27) and DefenseFinder v2.0.0 (28) with models v2.0.2 (**Dataset S1**). Predicted defense and anti-defense proteins were clustered across all 32,252 genomes using MMseqs2 (41) at 95% amino acid identity (**Dataset S2**). Results are reported in full in **Note S1**. Specific defense system repertoires of strains selected for RB-TnSeq library generation are listed in **Table S2**.

The phylogenetic tree of 32,252 O157:H7 genomes was constructed by the rapid neighbor-joining algorithm using genomic distances estimated by one-permutation hashing with optimal densification as implemented in bindashtree v0.1.2 (https://github.com/jianshu93/bindashtree) (42).

Outer surface glycan biosynthesis and outer membrane proteins relevant to phage infection were compared between strains ECRC98, ECRC100, ECRC101 and ECRC102 with parasail (43) to search for sequence polymorphisms.

### Phage susceptibility screening

Phage ability to infect the selected O157:H7 strains was tested in two successive screens. The first screen consisted in testing the ability of 161 diverse phages from the collection assembled by Moriniere et al. (12) to induce bacterial clearance at a single high titer. Lysates were normalized at approximately 10^9^ PFU/mL (titer calculated on the propagation host) and 2 µL droplets were spotted on bacterial lawns following the methodology described in **Bacteriophage strains and propagation**. Bacterial clearance was recorded manually after an overnight incubation at 37°C and scored as a binary phenotype. The phage gene-sharing network visualization was made with vContTACT 2.0 (44).

The second screen aimed at determining single plaque formation and efficiency of plating (EOP) of a subset of 38 phages on the 4 O157:H7 strains selected for further assays. Serial dilutions of phage lysates were prepared and spotted on bacterial lawns as described before. Single plaque formation was recorded as a binary qualitative metric (single plaque / no single plaque) after an overnight incubation at 37°C. Efficiency of plating was calculated as the ratio of the phage titer measured on the tested O157:H7 host vs. its titer measured on the propagation host. In the absence of visible single plaques, the highest dilution showing bacterial clearance was recorded and attributed a number of one plaque to allow EOP calculation and obtain a quantitative measurement of infection efficiency for every phage. Detailed results of both phage susceptibility screens were compiled in **Table S3**.

### Construction of *E. coli* O157:H7 RB-TnSeq libraries

Four *E. coli* O157:H7 RB-TnSeq transposon mutant libraries were constructed by conjugation using the diaminopimelic acid (DAP)-auxotrophic donor strain *E. coli* WM3064 carrying the pHLL250 barcoded mariner transposon vector library (strain AMD290) (13, 45, 46). The barcoded donor library was grown in LB supplemented with carbenicillin (50 μg/mL) and DAP (300 μM) at 37°C. The donor library culture was washed by centrifugation for 5 min at 7,168 rcf to remove carbenicillin and resuspended in 10 mL of fresh LB broth supplemented with DAP. Donor and recipient cultures were then mixed at a 1:1 ratio based on OD_600_ measurements and washed by centrifugating 5 min at 7,168 rcf. Appropriate controls were performed by mixing the donor and recipient cultures with LB broth. Cell pellets were resuspended in 1 mL of LB broth supplemented with DAP and deposited on LB agar plates to allow conjugation for 5 h at room temperature. Plates were then scrapped with fresh LB broth and the resulting slurry was mixed with glycerol at a 15% (v/v) concentration and stored at –80°C. Conjugation efficiency was estimated by selecting transconjugants cells on LB agar plates supplemented with kanamycin at 50 µg/mL. Frozen conjugation mix were subsequently diluted and plated in order to generate RB-TnSeq libraries containing at least 100,000 transconjugants. Colonies were scraped the next day with LB broth and pooled, and the suspension was diluted at a final OD_600_ of 0.5 and grown into 60 mL of fresh LB broth supplemented with kanamycin to purify the library from remaining intact donor vector. The resulting culture was then mixed with glycerol at a 15% (v/v) concentration, aliquoted in 1-mL cryotubes, and stored at –80°C. One aliquot per library generated was also immediately sacrificed to purify genomic DNA using the DNAeasy Blood and Tissue kit (QIAgen). Single-end (150bp) Illumina sequencing and mapping of random DNA barcodes to transposon insertion sites was conducted as described previously (47). The ECRC98 library contained 132,088 unique transposon mutants collectively covering 4,344 genes, the ECRC100 library 94,770 mutants covering 3,960 genes, the ECRC101 library 207,562 mutants covering 4,363 genes, and the ECRC102 library 184,092 mutants covering 3,997 genes.

### RB-TnSeq phage competitive fitness assays

Competitive fitness assays between the 4 RB-TnSeq libraries generated and the 38 phages selected were conducted in high-throughput liquid assays following the procedure established previously (12). Briefly, frozen aliquots of the libraries were thawed and grown in LB supplemented with kanamycin at 12.5 µg/mL until reaching mid-log phase. Library cultures were diluted to an OD_600_ of 0.04, dispensed into 48-well tissue culture plates, and mixed in equal volumes with phage lysates adjusted at 10^8^-10^9^ PFU/mL in a 700 µL final volume. Plates were sealed with Breathe-Easy® gas-permeable membranes (Sigma Aldritch) and incubated for 14 to 16 h with orbital shaking at 37°C in a BioTek 800 TS plate reader (Agilent Technologies). Surviving cells were collected, pelleted, and genomic DNA was extracted using the QIAamp 96 DNA QIAcube HT Kit (QIAgen).

Barcode amplification, BarSeq library preparation, and Illumina sequencing followed previously described protocols (48). BarSeq reads were computed to calculate gene fitness and t-test statistic scores with the FEBA code base (https://bitbucket.org/berkeleylab/feba/). The t-test statistic establishes the significance of the phenotype based on the consistency of the fitness signal in all the barcoded strains with unique transposon insertion sites located across the central 80% of the gene ORF (13). Consistency of the fitness signal across multiple mutants was also verified manually on an internal instance of the Fitness Browser (47) to resolve ambiguous fitness signals. Genes with a log_2_ fitness score at or above 4 and a t-test statistic at or above 5 were classified as high-confidence enriched loci. Genes with fitness scores at or below –2 and t-test statistic absolute value at or above 5 were classified as negative fitness loci.

### Experimental validation of gene fitness phenotypes in mutant strains

Targeted mutants were isolated directly from the RB-TnSeq library using a fitness-guided selection strategy in which phages associated with strong selective enrichment at defined loci were used to recover mutants from the pooled library. Individual mutants were identified using barcode-specific PCR with subsequent mapping to the disrupted locus.

Complementation plasmids were ordered from Twist Bioscience. Constructs carrying the native *ompA* promoter of strain ECRC100 (region defined as the 100 bp upstream of the start codon), the ORF of the gene of the interest, and its 100 bp downstream region, were cloned into the pTwist Chlor Medium Copy vector. The only exception was for the *tsx* complementation in ECRC100, for which the native 100 bp upstream and 100 bp downstream regions were used. The empty pTwist vector was generated by PCR amplification of the pTwist(*tsx)* plasmid using phosphorylated primers flanking the MCS in opposite directions. PCR amplification was carried out with Phusion™ high-fidelity DNA polymerase following manufacturer protocol. Self-ligation of the vector was carried out with T4 DNA ligase following manufacturer protocol, and the resulting product was transformed into *E. coli* DH5α chemically-competent cells (Thermo Fisher Scientific) following the manufacturer’s heat-shock protocol. Sequence was verified by whole plasmid sequencing at Angstrom Innovation using Oxford Nanopore Technology. Empty and complementation vectors were electroporated into the corresponding transposon mutant strains following standard procedures (49), and transformants were selected on LB agar supplemented with kanamycin and chloramphenicol.

Efficiency of plating assays were performed with selected bacterial mutant and phage strains to confirm the fitness signals yielded by pooled competitive fitness experiments. Experiments were conducted in biological triplicate following the same procedure for high-throughput phage titration described in **Bacteriophage strains and propagation**, but T-top agar was supplemented with CaCl_2_, kanamycin and chloramphenicol. Efficiency of plating was calculated as the ratio of PFU/mL measured on the mutant strain harboring the empty or complementation plasmid to PFU/mL measured on the wild-type host, and results are reported in **Dataset S12**.

## Data availability

Bacterial genome sequences generated in this study were deposited on NCBI GenBank under BioProject PRJNA1458855 with the following accession numbers: ECRC98 (*submitted*), ECRC100 (*submitted*), ECRC101 (*submitted*), ECRC102 (*submitted*). SRA accessions are listed in **Table S4**. Phage genome accessions and associated information are listed **Table S7**.

Defense system prediction results are available in **Datasets S1-S2**. Results of the initial phage susceptibility screen are computed in **Dataset S3**. Genes classified as having high-confidence positive or negative fitness scores in RB-TnSeq experiments are reported in **Datasets S4-S11**. Efficiency of plating experiment results are reported in **Dataset S12**. SNP-based clade assignments of 32,252 O157:H7 genomes are available in **Dataset S13**. All datasets, and supplementary **Tables S1-S7** and **Figures S1-S5**, are available at https://doi.org/10.6084/m9.figshare.33075194

## Acknowledgments

Authors would like to thank Morgan Price (LBNL) for processing all BarSeq and TnSeq runs. This work was supported by the National Science Foundation (NSF grant no. 2220735; EDGE CMT: Predicting bacteriophage susceptibility from *Escherichia coli* genotype), the U.S. Department of Agriculture National Institute of Food and Agriculture (USDA-NIFA, Hatch project no. PEN4826), and the National Center for Advancing Translational Sciences, National Institutes of Health (NIH grant no. TL1 TR002016).

