## Supplementary Figures for "Surface architecture of the bacterial envelope determines phage adsorption route in pathogenic *Escherichia coli* O157:H7"

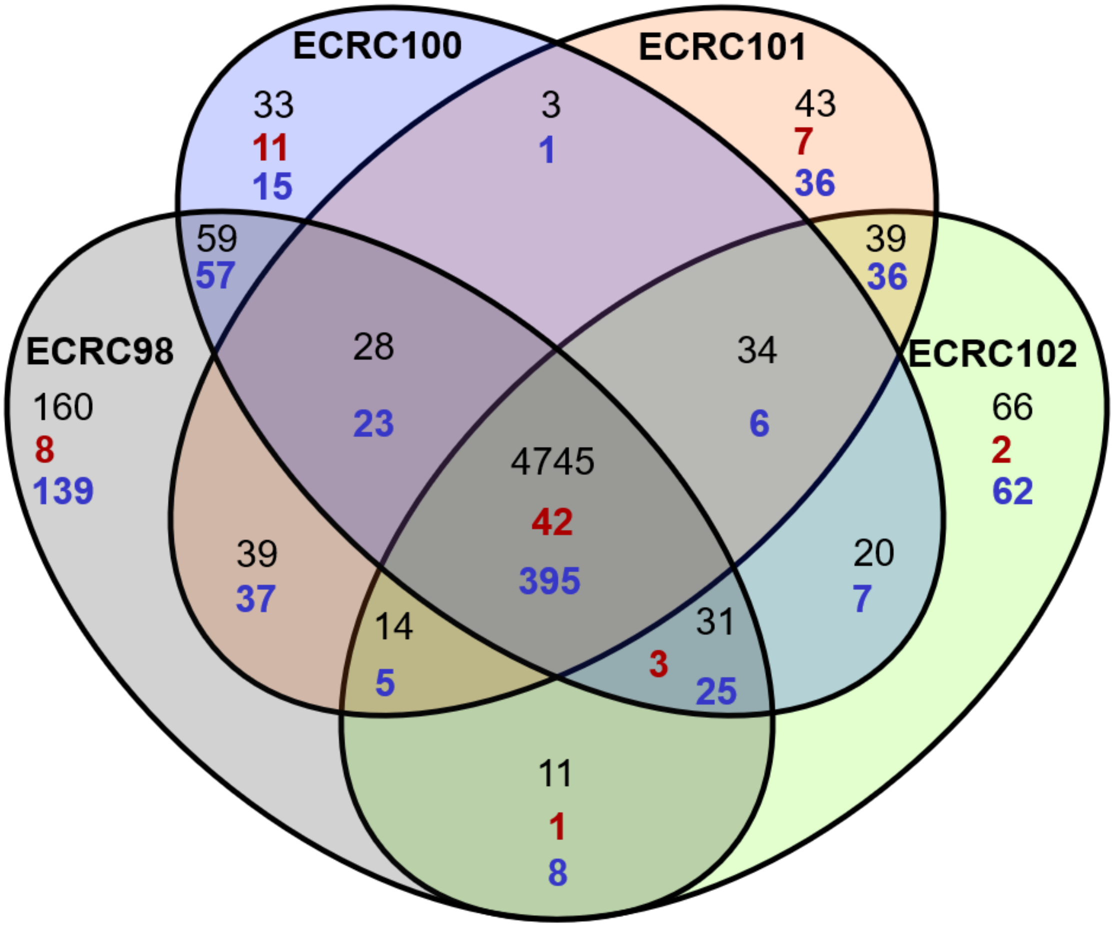


**Figure S1. Shared and strain-specific genes across the 4 O157:H7 strains selected**. Venn diagram representing the number of shared and unique genes between ECRC98, ECRC100, ECRC101, and ECRC102.

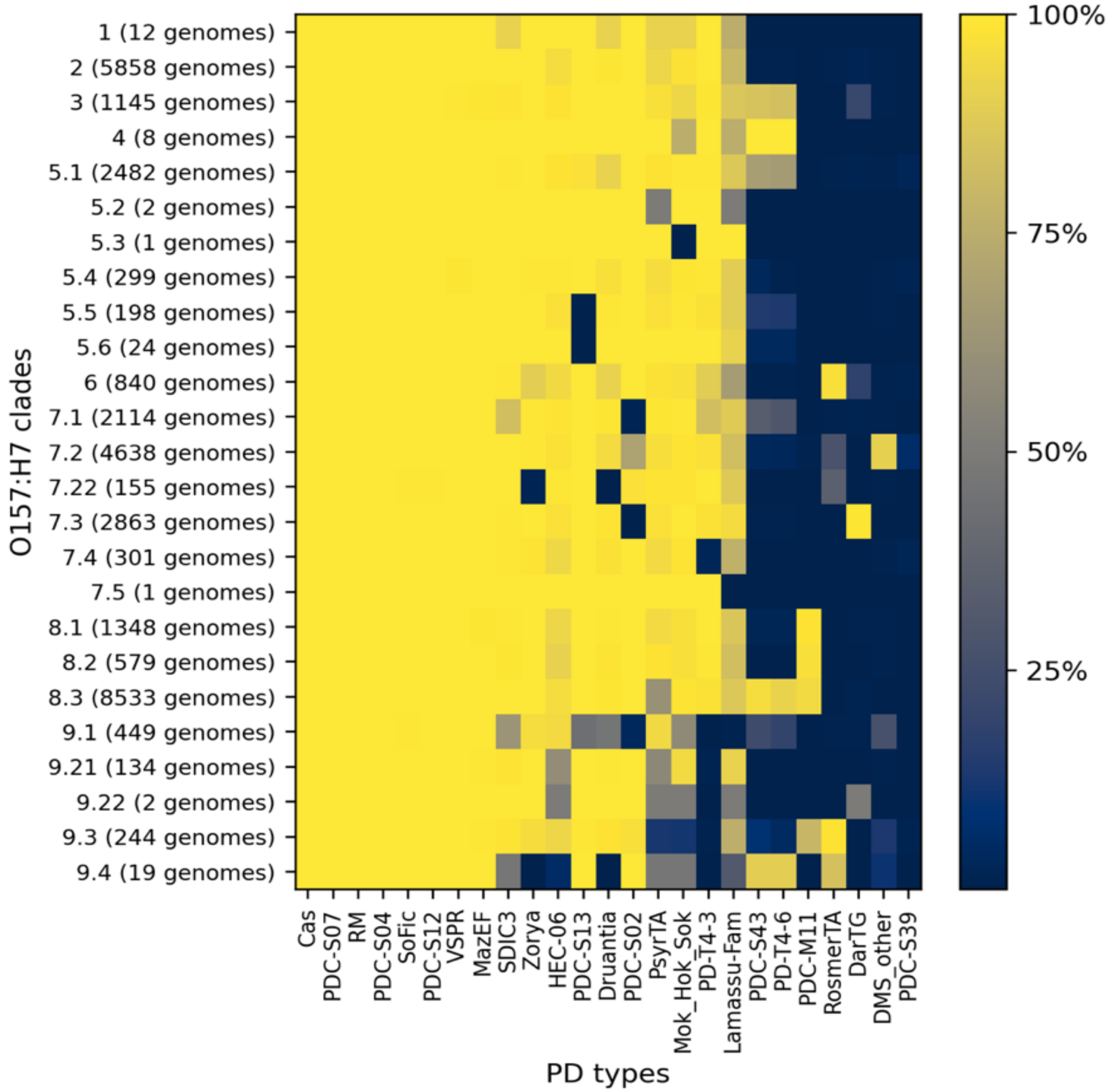


**Figure S2. Distribution of predicted prokaryotic defense systems across *Escherichia coli* O157:H7 clades.** Heatmap showing the prevalence of predicted defense system categories across O157:H7 clades. Rows represent phylogenetic clades (number of genomes in parentheses) and columns represent defense system types identified by PADLOC v2.0.0 and DefenseFinder v2.0.0 (models v2.0.2). Clades 9.3 and 9.4 correspond to *E. coli* O55:H7 lineages. A system was scored as present in a genome when at least one gene associated with that system was identified by either tool. Values represent the percentage of genomes within each clade encoding a given system.

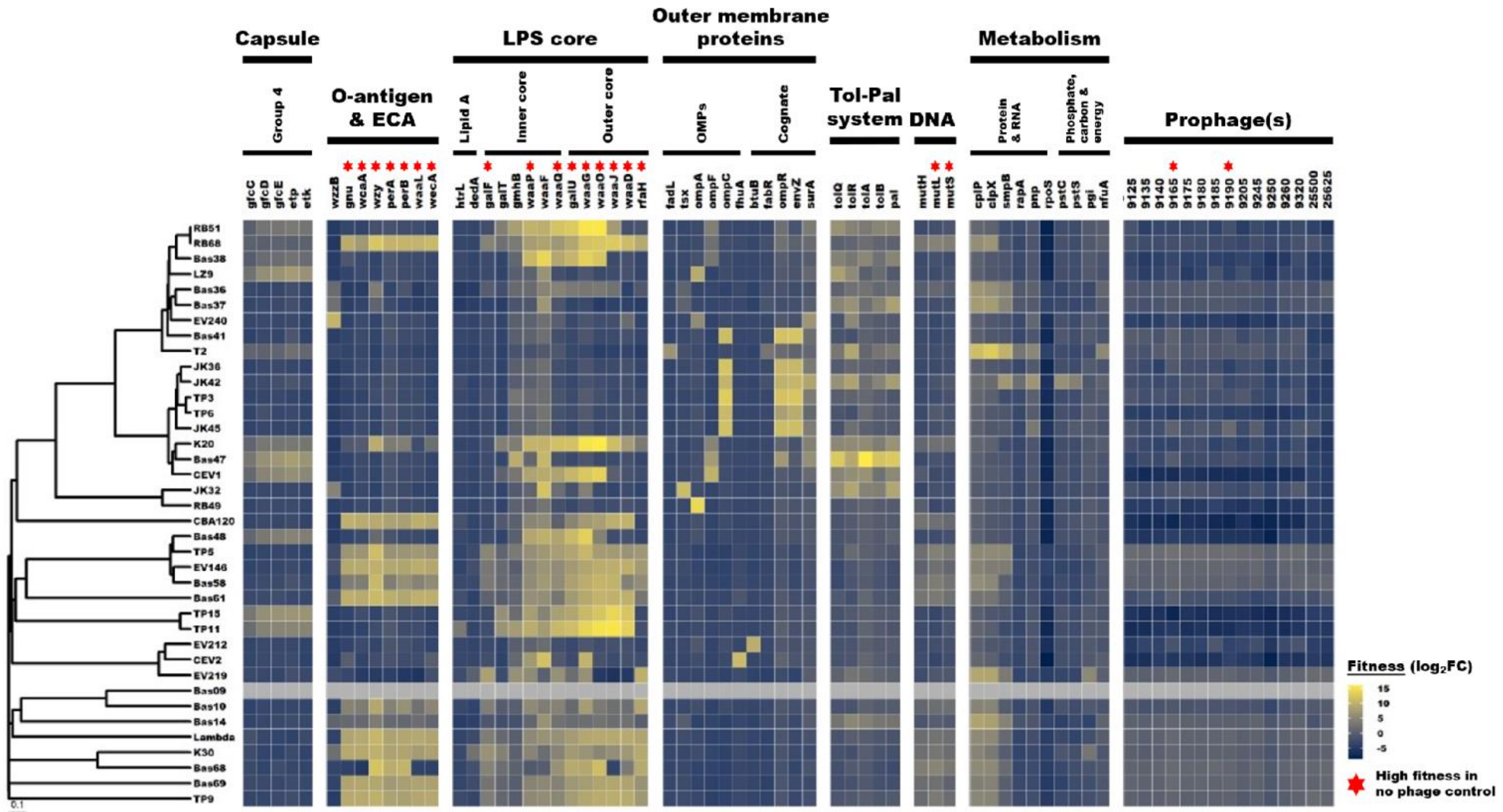


**Figure S3. Genome-wide identification of surface-associated determinants of phage susceptibility in *Escherichia coli* O157:H7 strain ECRC98.** Heatmap of loss-of-function (LOF) RB-TnSeq selected data for 37 dsDNA phages, where each row represents a phage and each column represents a gene. Fitness values are shown as $\log_2$ enrichment scores, where high values indicate a selective advantage under phage challenge, and low values indicates a detrimental fitness effect. Genes are grouped by functional categories, including group 4 capsule, lipopolysaccharide (LPS) core, O157 O-antigen, enterobacterial common antigen (ECA), outer membrane proteins (OMPs), and transport-associated functions. Other acronyms: RM = restriction-modification; RE = restriction endonuclease; MT = methyltransferase. Multi-gene signals were summarized into recognition of unique outer surface components (capsule, O157 O-antigen, R3-type LPS core sugars, OMPs). Phage dendrogram was computed based on pairwise proteomic equivalent quotient (PEQ) values.

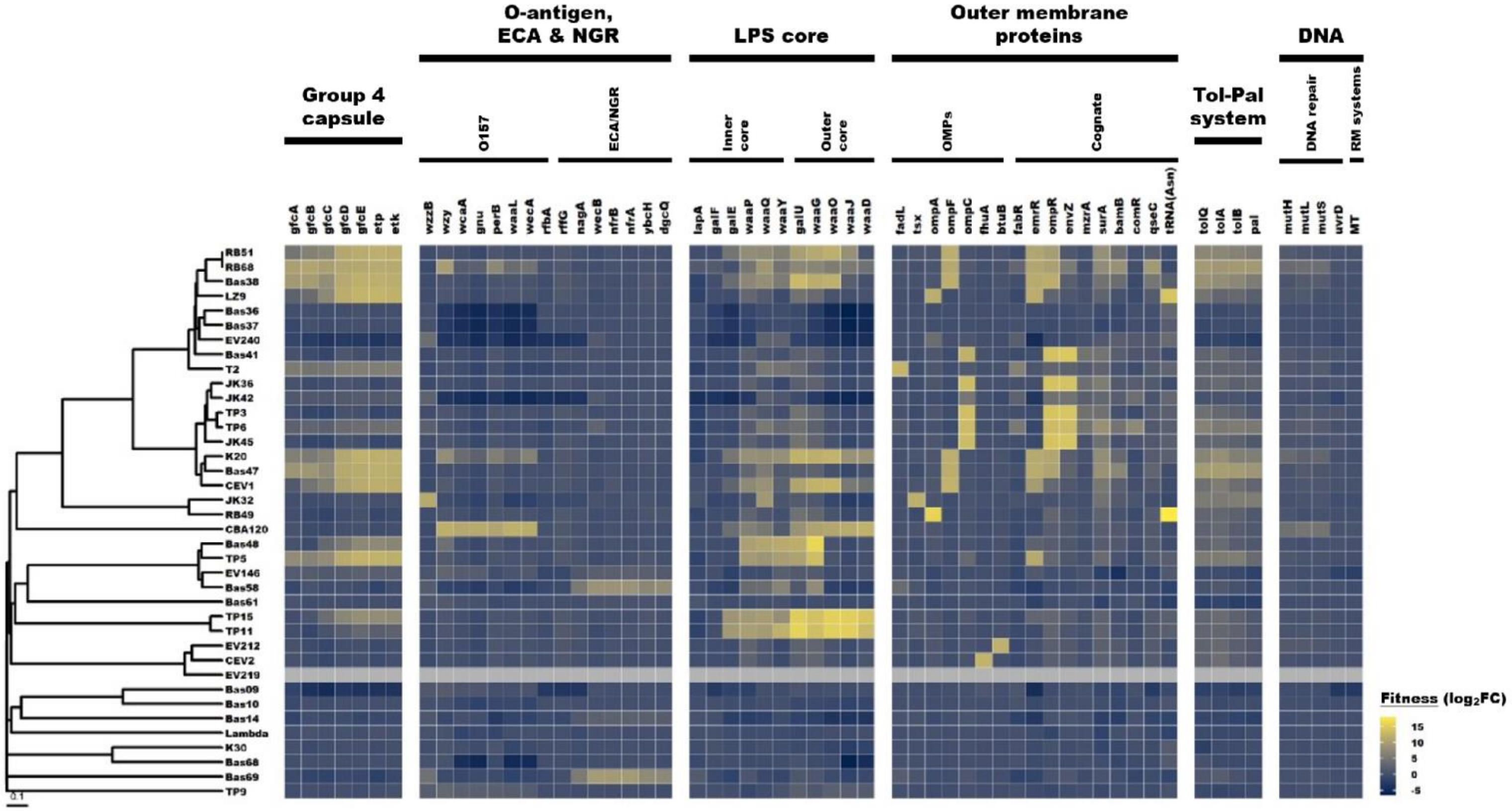


**Figure S4. Genome-wide identification of surface-associated determinants of phage susceptibility in *Escherichia coli* O157:H7 strain ECRC101.** Heatmap of loss-of-function (LOF) RB-TnSeq selected data for 37 dsDNA phages, where each row represents a phage and each column represents a gene. Fitness values are shown as $\log_2$ enrichment scores, where high values indicate a selective advantage under phage challenge, and low values indicates a detrimental fitness effect. Genes are grouped by functional categories, including group 4 capsule, lipopolysaccharide (LPS) core, O157 O-antigen, enterobacterial common antigen (ECA), outer membrane proteins (OMPs), and transport-associated functions. Other acronyms: RM = restriction-modification; RE = restriction endonuclease; MT = methyltransferase. Multi-gene signals were summarized into recognition of unique outer surface components (capsule, O157 O-antigen, R3-type LPS core sugars, OMPs). Phage dendrogram was computed based on pairwise proteomic equivalent quotient (PEQ) values.

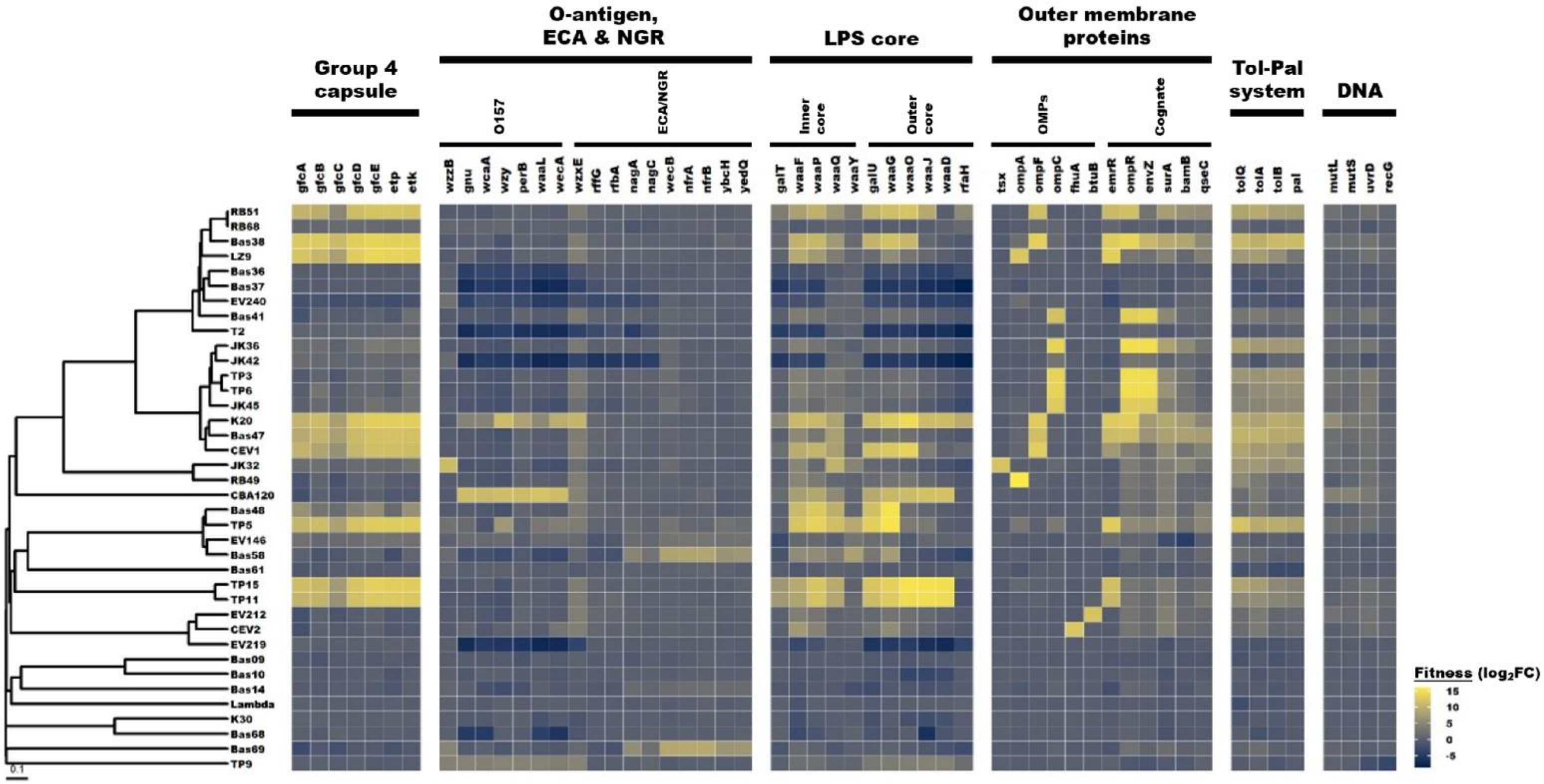


**Figure S5. Genome-wide identification of surface-associated determinants of phage susceptibility in *Escherichia coli* O157:H7 strain ECRC102.** Heatmap of loss-of-function (LOF) RB-TnSeq selected data for 38 dsDNA phages, where each row represents a phage and each column represents a gene. Fitness values are shown as $\log_2$ enrichment scores, where high values indicate a selective advantage under phage challenge, and low values indicates a detrimental fitness effect. Genes are grouped by functional categories, including group 4 capsule, lipopolysaccharide (LPS) core, O157 O-antigen, enterobacterial common antigen (ECA), outer membrane proteins (OMPs), and transport-associated functions. Other acronyms: RM = restriction-modification; RE = restriction endonuclease; MT = methyltransferase. Multi-gene signals were summarized into recognition of unique outer surface components (capsule, O157 O-antigen, R3-type LPS core sugars, OMPs). Phage dendrogram was computed based on pairwise proteomic equivalent quotient (PEQ) values.
